# *Yersinia pestis* actively inhibits the production of extracellular vesicles by human neutrophils

**DOI:** 10.1101/2024.12.20.629761

**Authors:** Katelyn R. Sheneman, Timothy D. Cummins, Michael L. Merchant, Joshua L. Hood, Silvia M. Uriarte, Matthew B. Lawrenz

**Affiliations:** Department of Microbiology and Immunology, University of Louisville; Department of Medicine and Proteomics Technology Center, University of Louisville; Department of Pharmacology and Toxicology, University of Louisville; Hepatobiology and Toxicology COBRE, University of Louisville; Department of Oral Immunology & Infectious Disease, University of Louisville; Center for Predictive Medicine for Biodefense and Emerging Infectious Diseases, University of Louisville

**Keywords:** *Yersinia pestis*, plague, type 3 secretion system (T3SS), Yop effectors, human neutrophils (hPMNs)

## Abstract

*Yersinia pestis* is the etiologic agent of the plague. A hallmark of plague is subversion of the host immune response by disrupting host signaling pathways required for inflammation. This non-inflammatory environment permits bacterial colonization and has been shown to be essential for disease manifestation. Previous work has shown that *Y. pestis* inhibits phagocytosis and degranulation by neutrophils. Manipulation of these key vesicular trafficking pathways suggests that *Y. pestis* influences EV secretion, cargo selection, trafficking, and/or maturation. Our goal was to define the EV population produced by neutrophils in response to *Y. pestis* and determine how these vesicles might influence inflammation. Toward these goals, EVs were isolated from human neutrophils infected with *Y. pestis* or a mutant lacking bacterial effector proteins known to manipulate host cell signaling. Mass spectrometry data revealed that cargoes packaged in EVs isolated from mutant infected cells were enriched with antimicrobials and cytotoxic proteins, contents which differed from uninfected and *Y. pestis* infected cells. Further, EVs produced in response to *Y. pestis* lacked inflammatory properties observed in those isolated from neutrophils responding to the mutant. Together, these data demonstrate that *Y. pestis* actively inhibits the production of antimicrobial EVs produced by neutrophils, likely contributing to immune evasion.

## Introduction

*Yersinia pestis* is the etiologic agent of the disease known as plague. This gram-negative bacterium can cause bubonic, pneumonic, or septicemic plague upon infection of the dermis, lungs, or bloodstream, respectively [1–3]. Person-to-person transmission is associated with pneumonic infection, by which the bacteria become aerosolized via exhalation. Without medical intervention, pneumonic infection can be lethal as early as 72 hours post-exposure [3]. A unique hallmark of pneumonic plague is the generation of a biphasic immune response [4–7]. During the first 36-48 h of infection, there is minimal inflammation at the site of infection, allowing *Y. pestis* to colonize the lungs unhindered from host intervention. However, by 48 h post-infection, there is a significant increase in inflammation involving the production of inflammatory mediators, such as TNF-α, IFN-γ, and leukotriene B4 (LTB_4_), as well as a robust influx of immune cell infiltrates [4, 8–10]. The generation and maintenance of this non-inflammatory environment is vital for colonization and disease progression [9].

While *Y. pestis* has a wide repertoire of virulence mechanisms, the Ysc type 3 secretion system (T3SS) and the effector proteins secreted through this system have a significant role in immune subversion [11, 12]. Upon translocation into host cells, the seven Yop effector proteins strategically manipulate host signaling pathways to suppress the generation of a productive immune response, resulting in the biphasic immune response characteristic of plague [4–7].

These associated effector proteins have been extensively studied for their ability to manipulate host signally pathways involved in inflammation and pathogen elimination [13]. Despite encoding only seven secreted effector proteins, these Yop effectors significantly inhibit the ability of host cells to respond to infection. For example, YpkA, YopE, YopH, and YopT systematically disrupt cytoskeletal actin dynamics by targeting Rho, Rac, and other focal adhesion proteins to inhibit many cellular responses [14–17], including vesicular trafficking and calcium signaling [18, 19]. YopJ disrupts MAPK signaling and NF-kB cascades that directly limit cytokine and chemokine production [19–22]. Finally, YopK and YopM strategically inhibit essential mediators of inflammasome activation and proinflammatory cell death pathways [23–27]. Together, the *Y. pestis* Yop effectors effectively inhibit the ability of the host to directly kill the pathogen and initiate a timely inflammatory response.

Polymorphonuclear neutrophils (PMNs) and macrophages are the initial cells that respond to *Y. pestis* during infection [7, 11]. As such, these are the primary cells targeted by the bacterium for Yop effector translocation during infection [4, 28, 29]. In a pneumonic model, PMNs represent >80% of the cell population intoxicated with Yop effectors by 12 h post-infection [7, 9], indicating that interactions with these cells is key to generating the early non-inflammatory environment needed to establish infection. In vitro studies have demonstrated that *Y. pestis* uses the Yop effectors to actively inhibit many of the antimicrobial mechanisms of the PMN, including phagocytosis, degranulation, and ROS production [13, 19]. Moreover, several of the Yop effectors inhibit the synthesis and release of inflammatory cytokines and lipids, effectively limiting recruitment of circulating immune cells into the tissue [10, 29, 30]. Together, these data indicate that manipulation of PMNs by *Y. pestis* is imperative for establishing lethal infection.

Extracellular vesicles (EVs) have become recognized for their preeminent role in mediating intercellular communication [31–33]. EVs are lipid-bound vesicles produced by a variety of cells, including immune cells. EVs can be produced by two primary pathways. Small EVs (10-200nm in diameter, historically referred to as exosomes) are produced within the multivesicular body, where proteins, lipids, and nucleic acids are strategically packaged prior to release via exocytosis [34–36]. Large EVs (>200nm in diameter, historically referred to as microvesicles) are produced via plasma membrane budding and contain cellular mediators abundant within the cytoplasm [37]. Once released, EVs can interact with other cells to establish biochemical communication. Importantly, EV payloads change in response to the physiological environment of the cell, and these changes dictate the signaling potential of EVs [32, 38, 39]. In the context of infection, EVs produced by sentinal leukocytes relay pro-inflammatory signals and/or PAMPs that promote the mobilization of naïve immune cells [32, 40]. Moreover, EVs can augment macrophage polarization, induce TLR signaling, and drive immune cell chemotaxis [41–44].

Additionally, EVs can relay these signals in a paracrine and endocrine manner, highlighting their role in intercellular communication and immune stimulation during infection [43, 45, 46]. While growing data supports that EV production and payloads are key mediators needed for a timely and proper immune response, the EV response by leukocytes during plague has not been previously investigated. Here, we show for the first time that *Y. pestis* actively manipulates EV production by human PMNs, significantly impacting downstream EV function and signaling potential.

## Methods

### Isolation of PMNs and Macrophages

Human PMNs (hPMNs) and monocytes were isolated from venous blood using Ficoll density gradient separation as previously described [47]. Written consent was procured from each donor volunteer in accordance with the Institutional Review Board at the University of Louisville (IRB number 96.0191). For hPMNs, all preparations were >92% pure and utilized within 1 h of isolation. Peripheral human monocytes were differentiated into macrophages (hMDMs) by serum limitation as previously described [48, 49]. Briefly, monocytes were first cultured in RPMI supplemented with 20% fetal bovine serum (FBS). After 3 days, the medium was removed and replaced with RPMI + 10% FBS. After 2 days, the medium was replaced again with RPMI + 5% FBS. Finally, after 1 day, the medium was replaced with RPMI + 1% FBS. hMDMs were then used for subsequent studies on day 8.

### Bacterial and Cell Culture

All studies were performed with *Y. pestis* KIM1001 pgm(-) derivatives, which lack the chromosomally-encoded high pathogenicity pgm locus, allowing for experimentation at biosafety level 2 [50]. Bacteria were cultivated with aeration for 15-18 h in Brain Heart Infusion (BHI) broth (Difco BHI, Becton Dickinson) at 26°C. Prior to cell infection studies, bacteria were diluted 1:10 in fresh BHI supplemented with 20 mM MgCl_2_ and 20 mM Na-oxalate and grown at 37°C for 3 h.

### EV Isolation

EVs were isolated from hPMNs as previously described [40]. Briefly, hPMNs in suspension were incubated with *Y. pestis* at an MOI of 50 with rocking at 37°C. At 1 h, cells were incubated on ice for 10 min followed by centrifugation at 4,000xg for 20 minutes at 4°C to remove hPMNs.

Supernatants were passed through a 0.45µm CA filter (VWR, Cat. No. 76479-040) to further remove bacteria and large cellular debris. EVs were then isolated and concentrated by ultracentrifugation at 160,000xg for 55 minutes. The supernatant was removed and EVs were resuspended in 100μL of 1x PBS and stored at 4°C or -80°C.

### Characterization of EVs

Dynamic light scattering (DLS) was performed on the DynaPro plate reader (Wyatt Technologies), parameters set for an acquisition number of 20 and auto-attenuation time of 60 seconds. Autocorrelation for each acquisition was analyzed via regularization fit and size distribution analyzed by percent intensity. Nanoparticle tracking analysis (NTA) was performed

by Alpha Nano Tech, using a Zetaview Quatt (particle Metrix) instrument, equipped with a 488nm laser and sCMOS camera. Protein quantification was performed using Protelite Fluorometric Protein Quantification Kit (ThermoFisher, Cat. No. Q33211) optimized for the Qubit fluorometer.

### Transmission Electron Microscopy (TEM)

EVs were prepared for TEM as described previously [51]. Briefly, EVs were placed onto cross-hatched nickel grids at room temperature for 20 minutes. Grids were sequentially washed 2x with PBS and 2x with deionized water. After washing, grids were negatively stained with 0.3% uranyl-acetate for 5 minutes and subsequently washed 3x with deionized water. Images were collected via Hitachi HT7700 transmission electron microscope.

### Liquid Chromatography with Tandem Mass Spectrometry (LC-MS-MS)

LC-MS-MS to identify proteins associated with purified EVs was performed as previously described [52, 53]. Briefly, ∼50μg of each sample was vacuum dried and resuspended in 5% SDS, 50 mM triethyl ammonium bicarbonate (TEAB) in a standard S-trap (suspension trapping) proteomic workflow. Samples were vortexed and centrifuged prior to reduction in 25mM tris (2- carboxymethyl)phosphine (TCEP) at 65C for 30 minutes and cooled then alkylated with 20mM iodoacetamide (IAA) at room temperature for 20 minutes. Samples were acidified with 1.2% phosphoric acid and subsequently digested in 50mM TEAB with 80 ng/μg of trypsin for 2 hours at 47C without shaking. Peptides were eluted and vacuum dried then resuspended in 0.1% formic acid and diluted to 200ng/µL. Equal mass of peptides (∼600ng) were injected into a one-dimensional reverse phase liquid chromatography column and eluted into an Orbitrap mass spectrometer for MS/MS spectral acquisition. Spectral raw files were submitted to PEAKS X Studio for peptide and protein assignment using a Human FASTA database (version 20240425) for *in silico* mapping of peptides and proteins. Spectral matches were processed using Scaffold (version Q+ Scaffold_5, Proteome Software Inc, Portland,). Protein identifications were accepted if they could be established at greater than 99% probability to yield a false discovery rate less that 1.0%. Comparative protein analysis was performed using MetaboAnalyst by comparing spectral counts normalized to the mean [54]. Subsequent pathway analysis was performed using the Database for Annotation, Visualization, and Integrated Discovery (DAVID) [55].

### Bacterial Survival Assay

100 µL of EVs (derived from 1×10^8^ hPMNs) were added to 5×10^7^ opsonized bacteria in Hank’s buffered salt solution (HBSS) and incubated at 37°C for 40 minutes. Following the incubation, 2 mL of cold 1 mg/mL saponin in HBSS was added to lyse EVs and samples immediately frozen at -80°C for 20 minutes. Samples were thawed, serially diluted on BHI plates, and incubated at 26°C for 2 days to calculate CFU [39].

### hMDM Polarization and Flow Cytometry

Differentiated hMDMs were treated with LPS (MilliporeSigma, Cat. No. L2880) and IFNγ (Cell Signaling, Cat. No. 80385S) or EVs for 24h at 37°C and 5% CO_2_. Cells were removed from the plate, centrifuged at 6,000xg for 1 min, and fixed in 1% PFA on ice for 20 minutes. Cells were washed two times with 1x PBS and permeabilized with 0.5% Triton X-100 at room temperature for 15 min. Permeabilized cells were washed two times with 1x PBS + 10% BSA prior to staining. For staining, cells were incubated with Human TruStain FcX Blocking Solution (Biolegend, Cat. No. 422301) followed by anti-CD68 (VWR, Cat. No. 76322-418) and anti-CD80 (Thermo, Cat. No. 14292-AP0) antibodies for 45 min at 4°C. Cells were pelleted and resuspended in PBS. Single cell suspensions were generated by filtering through 70um mesh prior to analysis. Mature macrophages were identified as cells with high expression of CD68. M1-polarized macrophages were identified as cells with high expression of both CD68 and CD80.

### Bacterial Survival in Macrophages

5×10^5^ hMDMs were transferred into individual wells of a 96-well plate and treated with 100 µL of EVs. After 24h, cells were washed with serum-free RPMI and incubated at 37°C with 5×10^6^ bacteria. 20 min post-infection, hMDMs were treated with 8 µg/mL gentamicin to eliminate extracellular bacteria. One hour after gentamicin treatment, the supernatant was removed and replaced with medium containing 2 µg/mL gentamicin. Bacterial survival was assessed at 6 h post-infection via conventional CFU enumeration [49].

## Results

### hPMN EV characteristics change in response to *Y. pestis*

*Y. pestis* inhibits both endocytic and exocytic vesicular trafficking by hPMNs via the activity of the Yop effectors [10, 19, 29, 50], suggesting that vesicular trafficking pathways leading to EV production by hPMNs may also be altered by *Y. pestis*. To test this hypothesis, EVs were isolated by differential centrifugation from uninfected hPMNs (UI) or hPMNs following 1 h infection with *Y. pestis* lacking the pgm locus (Yp; *Y. pestis* inhibition of vesicular trafficking is independent of the factors encoded by the pgm locus) or Yp lacking the seven Yop effector proteins (T3E) [50]. Nanoparticle tracking analysis indicated no significant differences in the total number of EVs produced by UI and Yp infected hPMNs, but significantly higher concentrations of EVs were isolated from hPMNs infected with T3E (Figure 1a, p<0.001). As expected for EVs, the particles were sensitive to Triton X-100 treatment (Figure 1b). While *Y. pestis* can produce outer membrane vesicles (OMVs), we were unable to isolate detectable levels of OMVs from exclusively bacterial cultures under these conditions, supporting that those vesicles isolated were primarily derived from hPMNs (Figure S1). TEM imaging further confirmed the presence of semi-round, cup-shaped vesicles, consistent with the morphology typically observed for EVs (Figure 1c) [56]. The mean EV diameter isolated from UI hPMNs was 210.7±19.07 (Figure 1d), which was significantly larger than the mean diameter of EVs produced in response Yp or T3E infection (149±8.83 and 149±7.73, respectively; p<0.0001).

**Figure 1:**
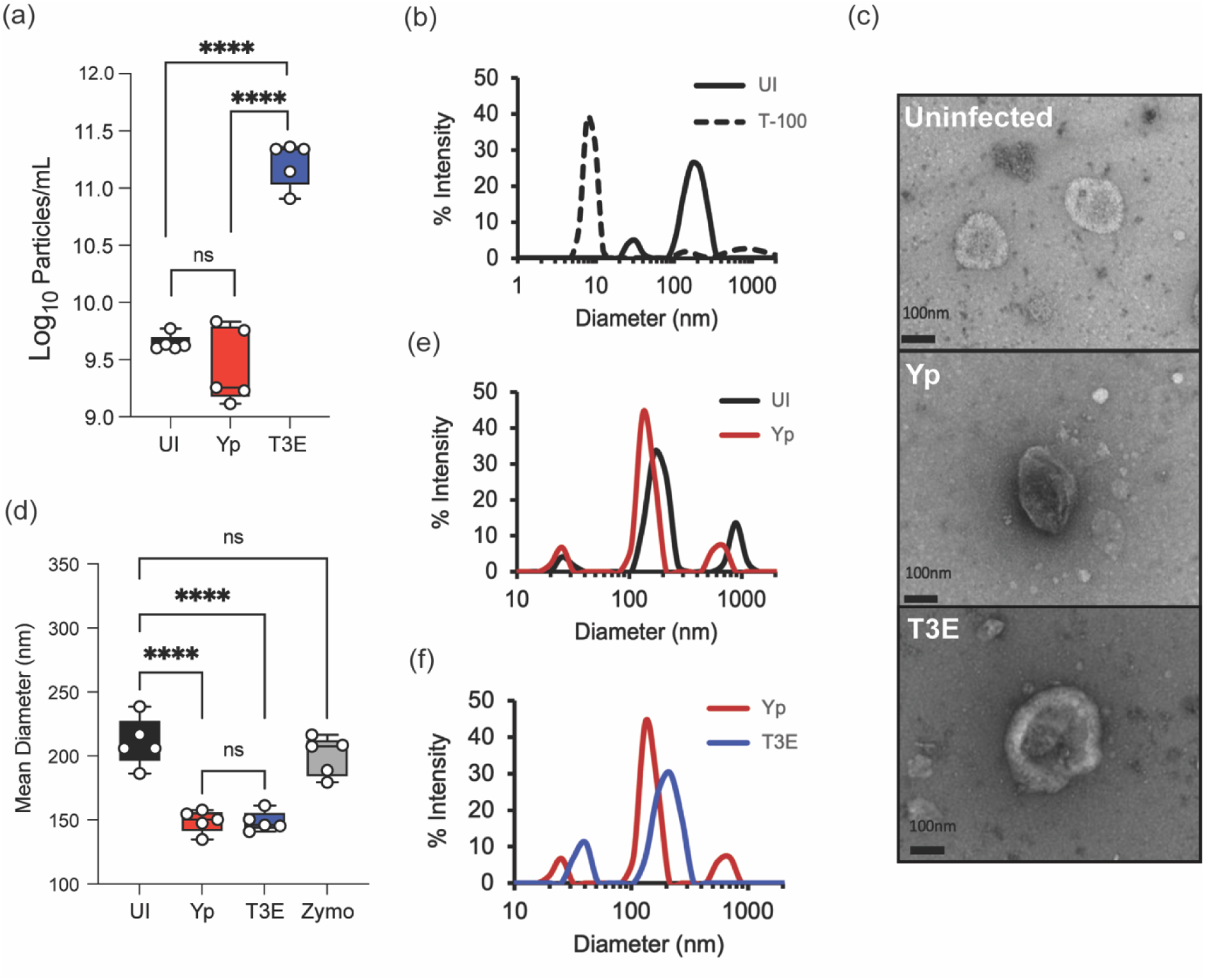
**Characterization of *Y. pestis*-elicited EVs.** hPMNs were infected with *Y. pestis* (Yp; red) or *Y. pestis* lacking the Yop effectors (T3E; blue) for 1 hour prior to EV isolation via ultracentrifugation. (a) Total EV particle quantification via NTA. Each point represents EVs from an individual human donor. (b) Representative DLS analysis of EVs from uninfected hPMNs without (UI; solid black line) or with Triton-X100 treatment (T-100; dashed line). (c) Representative TEM images of EVs isolated from uninfected, Yp, or T3E-infected hPMNs. Original magnification ∼40,000x. (d) Average diameter of EVs isolated from uninfected, zymosan-treated (Zymo), Yp-infected, or T3E-infected hPMNs. Each point represents EVs from an individual human donor. (e-f) Representative DLS analysis of EVs isolated from UI, Yp, or T3E-infected hPMNs. (a,d) One-way ANOVA with Tukey; ns=not significant; ****=p≤0.0001. (b,e,f) Representative results of 5 independent donors.

EVs elicited by Yp and T3E infected hPMNs were also smaller than those isolated from zymosan-stimulated hPMNs (Figure 1d) [39]. However, despite similarities between mean diameters, dynamic light scattering revealed distinct profiles of EVs from the Yp or T3E infected cells, which also differed from UI hPMNs (Figures 1e and 1f). Together, these data suggest that EV populations produced by hPMNs change in response to *Y. pestis*, but also that the Yop effector proteins impact EV production during infection.

### *Y. pestis* infection alters which proteins are packaged into EVs by hPMNs

Given the fundamental role of EVs in mediating cellular communication during infection, we sought to explore the implications of *Y. pestis* infection on EV biogenesis, specifically discrepancies in protein packaging. While the total protein concentration of the isolated EVs was comparable between UI and Yp samples, EVs isolated from T3E-infected hPMNs exhibited consistently higher protein yield (Figure 2a; p<0.0001). While this increased protein concentration could reflect increased EV production (Figure 1a), it also suggested changes in protein packaging within the EV population. To determine if *Y. pestis* alters which proteins are packaged into EVs, the proteomic profile of EVs isolated from hPMNs was determined by high resolution mass spectrometry. Regardless of source, all isolated EVs were enriched with proteins recognized as EV markers as put forth by MISEV2023 (minimal information for studies of extracellular vesicles 2023; [56]), confirming the abundance of EVs within our samples (Figure 2b). However, strikingly distinct proteomic profiles were observed depending on hPMN treatment (Figures 2c-e). Dimensional reduction using PLS-DA for comparison revealed that each condition displayed unique proteomic composition and that these divergencies were consistently reproducible (Figures 2c and 2d). Despite the similarity in protein concentrations, the protein composition of EVs isolated from Yp-infected hPMNs were starkly different from UI hPMNs (Figures 2c-e). Moreover, there were also significant differences in the proteins packaged into the EVs isolated from T3E-infected hPMNs compared to either of the other two conditions. The disparate enrichment of the proteins in each of these groups suggests that *Y. pestis* strategically manipulates the packaging of proteins within EVs.

**Figure 2:**
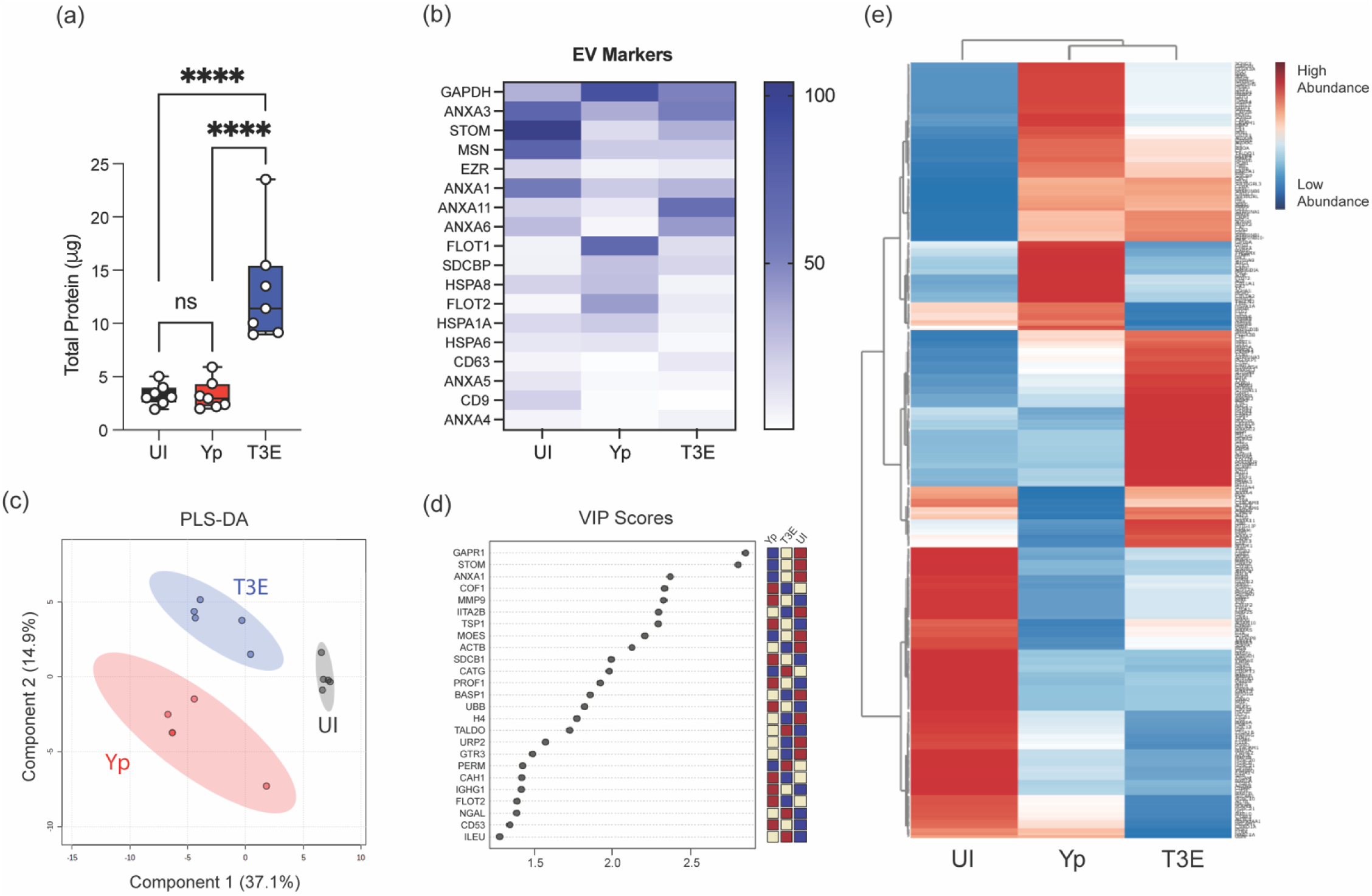
**Protein content of EVs is altered by *Y. pestis.*** hPMNs were infected with *Y. pestis* (Yp; red) or *Y. pestis* lacking the Yop effectors (T3E; blue) for 1 hour prior to EV isolation via ultracentrifugation. (a) Protein quantification of indicated EVs. Each point represents EVs from an individual human donor. One-way ANOVA with Tukey; ns=not significant; ****=p≤0.0001. (b) Average enrichment of recognized EV markers as measured by MS (n=5). (c) Partial Least Squares Discriminant Analysis (PLS-DA) plot depicting discriminant analysis of EVs isolated from uninfected (UI), Yp, or T3E-infected hPMNs. (d) VIP scores contributing to the variance in (c). (e) Heat map depicting protein enrichment EVs isolated from uninfected (UI), Yp, or T3E-infected hPMNs (average from 5 individual donors for each group). Proteins clustered according to Ward’s Hierarchical Agglomerative Clustering Method [54].

To further characterize the EV proteomes, the identified proteins were subcategorized based on statistical differences in enrichment compared to EVs from UI cells (Figure 3a, p<0.05). For the Yp-elicited EVs, 66 proteins were significantly enriched and 97 significantly deficient compared to UI EVs. A similar number of proteins were found dysregulated for the T3E EVs (68 proteins enriched; 95 proteins deficient), but the majority of these proteins were not conserved between two infection conditions. From this analysis, we identified three trends in differential protein enrichment: 1) proteins enriched in UI EVs but subsequently diminished in both Yp and T3E groups, 2) proteins overrepresented in EVs in response to *Y. pestis* infection regardless of bacterial strain, and 3) proteins enriched in T3E-elicited EVs but significantly diminished in Yp- elicited EVs.

**Figure 3:**
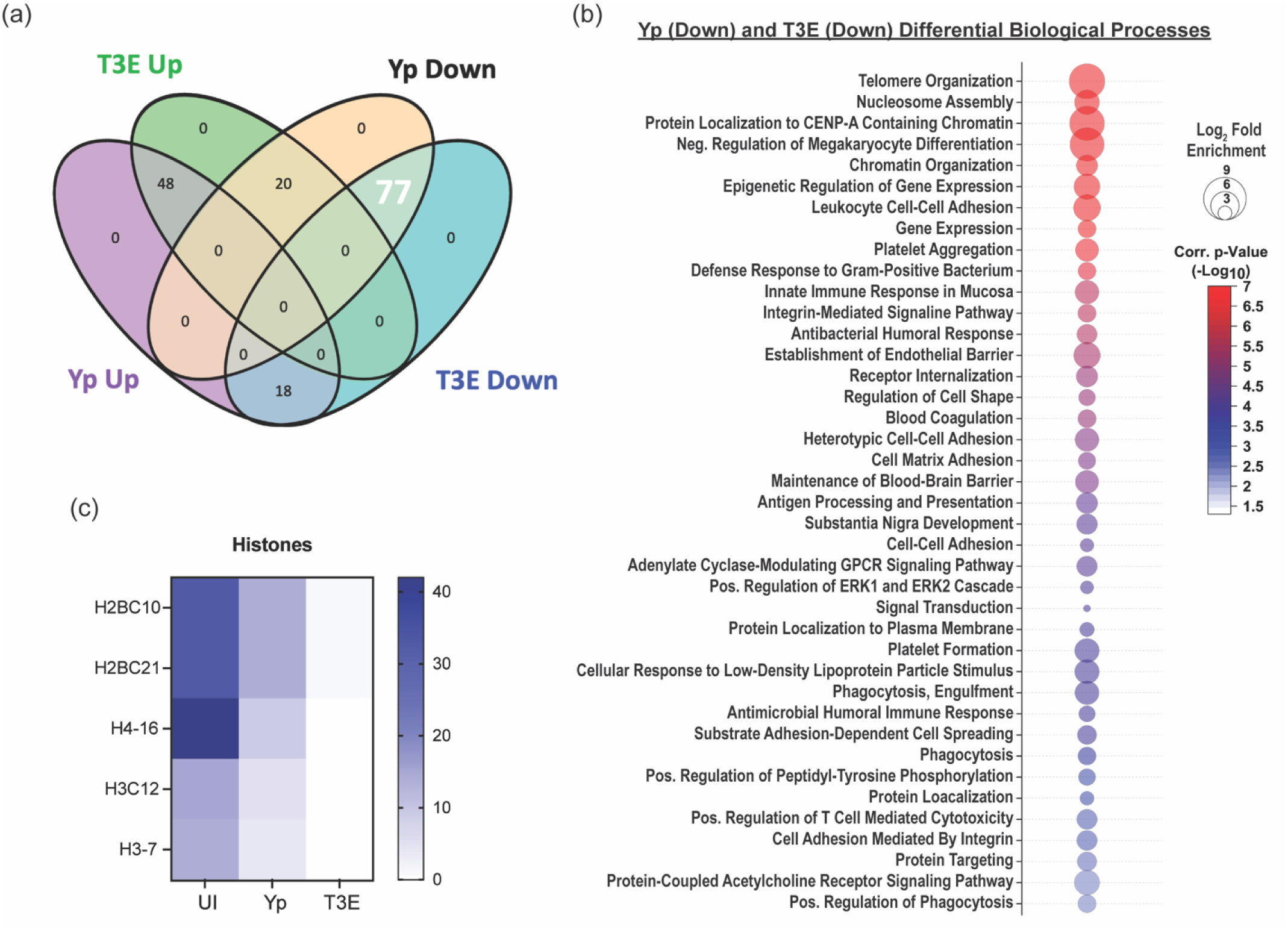
**EV proteins reduced in response to *Y. pestis.*** (a) Venn diagram highlighting the 77 proteins (white) that were enriched in EVs from UI hPMNs (p<0.05). (b) Biological process analysis predicted by DAVID [55] using the 77 enriched proteins (pathway enrichment displayed -Log_10_ p>2.5). (c) Prevalence of histone proteins from EVs isolated from UI, Yp-infected, or T3E-infected EVs.

The largest group of changes we observed were 77 proteins that were reduced in EVs isolated from infected cells compared to UI cells (Figure 3a). Pathway analysis revealed a diverse repertoire in the biological processes of the proteins packaged in response to infection (Figure 3b). The most significant changes were in proteins associated with nucleosome assembly and DNA binding and were highlighted by significant changes in the packaging of numerous histone proteins (Figure 3c). The second major trend was enrichment of 48 proteins in EVs isolated after infection regardless of the bacterial strains (Figure 4a). This included enrichment for proteins affiliated with extracellular matrix (ECM) remodeling as well as regulation of inflammatory processes (Figure 4b), highlighted by increased packaging of several ECM proteases and nutritional immunity proteins (Figures 4c and 4d). Lastly, we identified 20 proteins highly enriched in T3E-elicited EVs but significantly less prevalent in Yp-elicited EVs relative to EVs from uninfected cells (Figure 5a), the majority of which were associated with host defense and immune regulation (Figure 5b). Of these, we observed a significant absence of Annexin proteins in Yp-elicited EVs, a group of proteins associated with EV biogenesis (Figures 5c and 5d). Moreover, we observed an enrichment for 10 proteins with direct antimicrobial activity (e.g. MPO, CTSG, DEFA1B), which also represented the most disparately packaged proteins between Yp and T3E EVs (Figure 5e), suggesting that the *Y. pestis* T3SS actively inhibits the packaging of antimicrobials that could significantly impact hPMN responses during plague.

**Figure 4:**
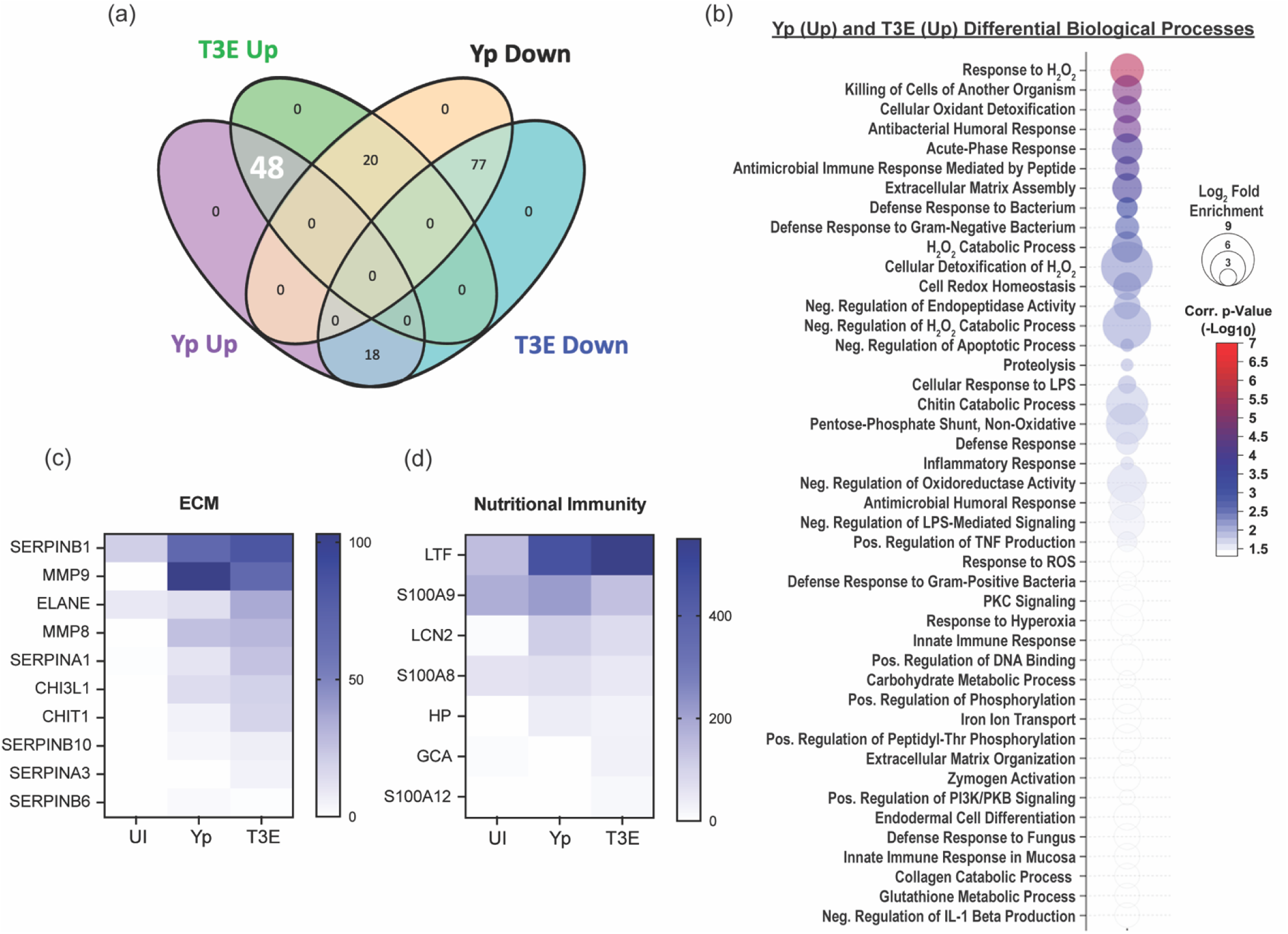
**EV proteins enriched in response to *Y. pestis.*** (a) Venn diagram highlighting the 48 proteins (white) that were enriched in EVs from infected hPMNs (p<0.05) compared to the UI EV population. (b) Biological process analysis predicted by DAVID [55] using the 48 enriched proteins (pathway enrichment displayed -Log_10_ p>1.5). Prevalence of ECM proteins (c) and nutritional immunity proteins (d) from EVs isolated from UI, Yp-infected, or T3E-infected EVs.

**Figure 5:**
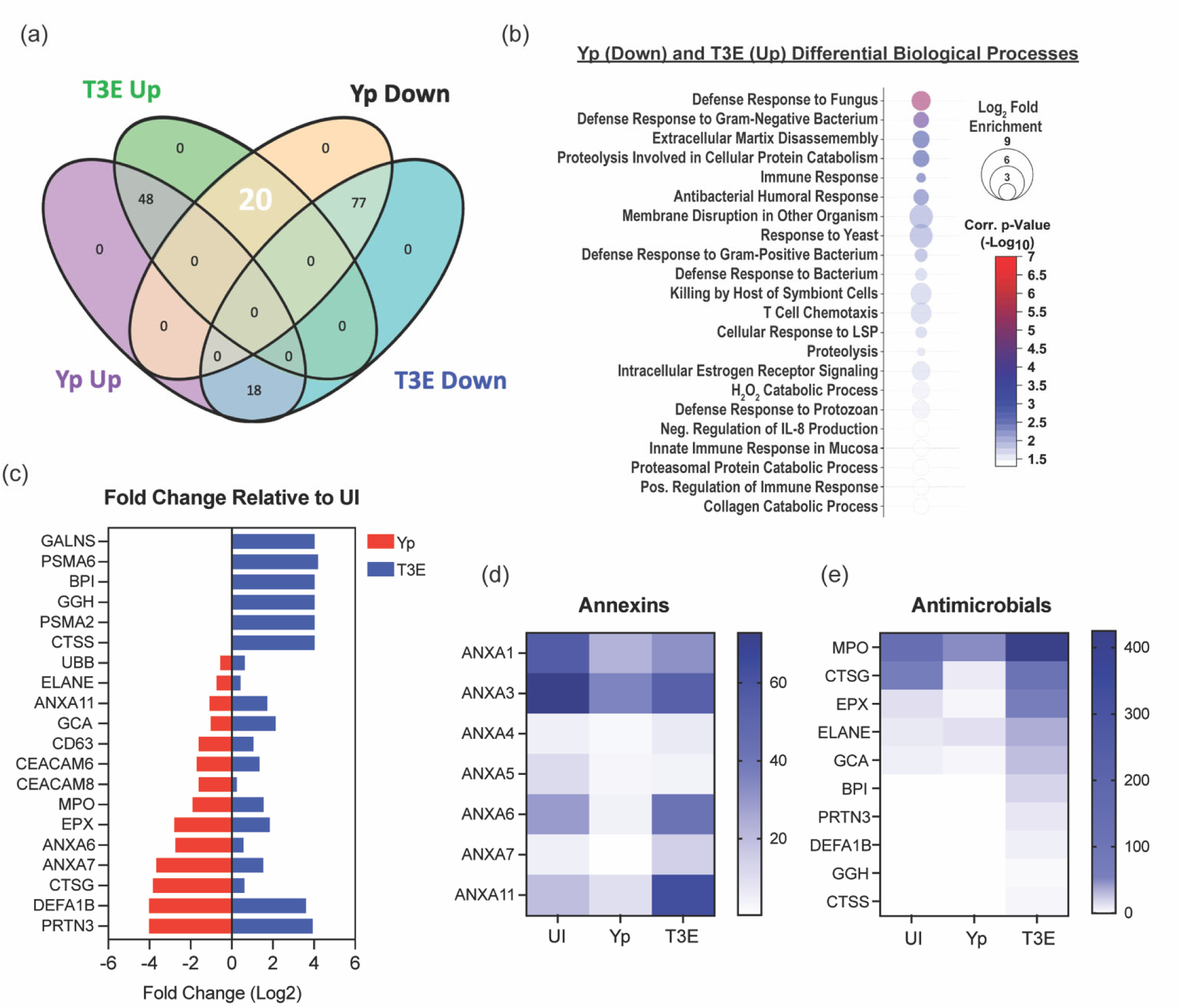
***Y. pestis* T3SS limits antimicrobial and pro-inflammatory protein packaging within EVs.** (a) Venn diagram highlighting the 20 proteins (white) that were enriched in T3E EVs and subsequently reduced in Yp EVs (p<0.05) compared to the UI EV population. (b) Biological process analysis predicted by DAVID [55] using the 20 dysregulated proteins (pathway enrichment displayed -Log_10_ p>1.5). (c) Fold change of the 20 identified proteins relative to UI. Prevalence of Annexins (d) and antimicrobial proteins (e) from EVs isolated from UI, Yp-infected, or T3E-infected EVs.

### *Y. pestis* inhibits the antimicrobial capacity of hPMN-derived EVs

Previous studies have demonstrated that EVs produced by activated hPMNs in response to other bacteria can have direct antimicrobial potential [40, 57, 58]. Given that we observed distinct differences in the antimicrobial proteome of EVs elicited by hPMNs in response to Yp and T3E (Figure 5e), we next sought to assess the antimicrobial capacity of EVs isolated from the infected hPMNs. While treatment of *Y. pestis* with EVs from UI hPMNs had a negligible impact on bacterial survival, treatment with EVs isolated from Yp-infected hPMNs seemed to slightly increase bacterial recovery (Figure 6a; p≤0.05). However, treatment with EVs isolated from T3E-infected hPMNs resulted in significantly lower bacteria recovery (p≤0.001), supporting that the enrichment of antimicrobial proteins in the T3E-elicited EVs potentiate direct antimicrobial activity.

**Figure 6:**
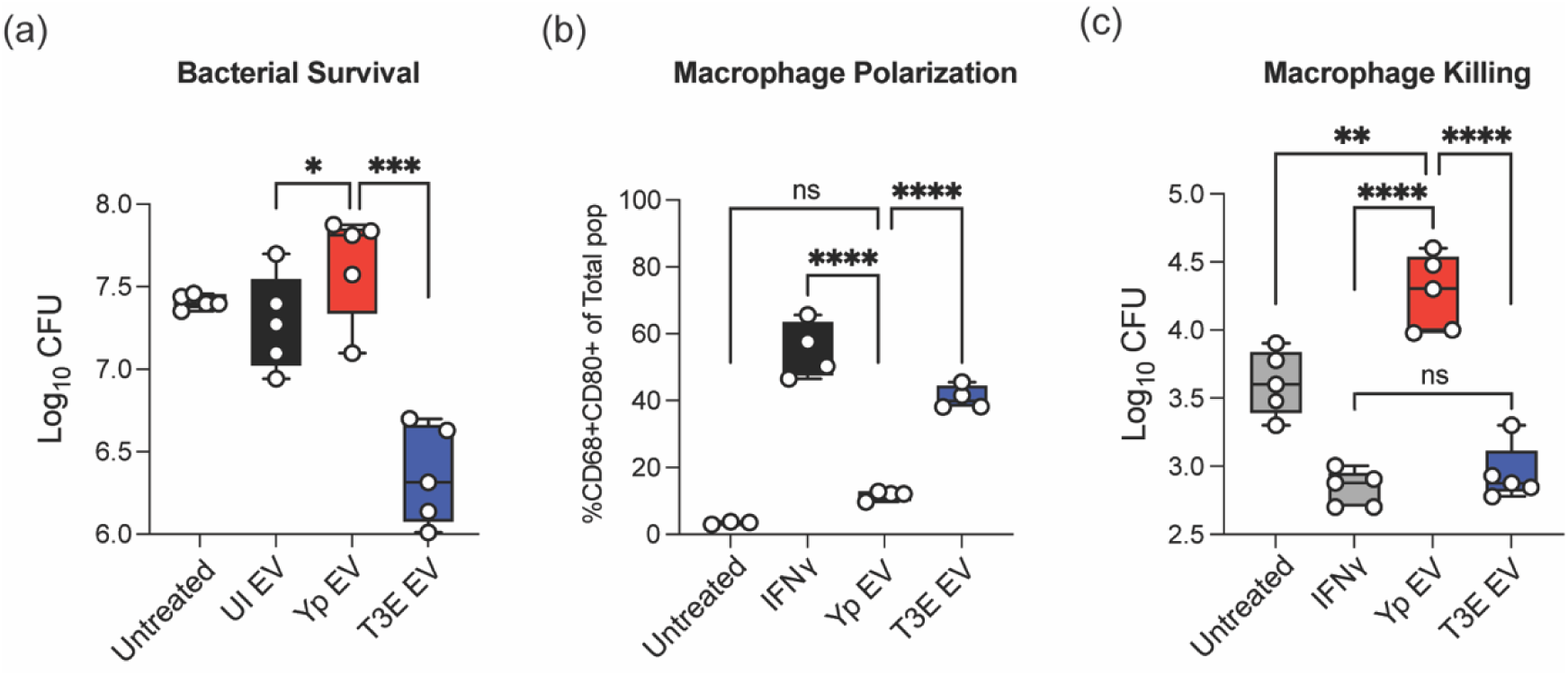
**Yp-elicited EVs have limited antimicrobial capacity.** (a) *Y. pestis* was treated for 40 min with EVs isolated from UI, Yp-infected, or T3E-infected hPMNs and bacterial viability was determined by serial dilution and conventional colony forming unit (CFU) formation on agar plates. (b) hMDMs were treated with EVs isolated from UI, Yp- infected, or T3E-infected hPMNs or IFNγ for 24 h and M1 polarization was determined by flow cytometry. (c) hMDMs were treated with EVs isolated from UI, Yp-infected, or T3E-infected hPMNs or IFNγ for 24 h and subsequently infected with Yp at an MOI of 10. Intracellular bacterial survival was assessed with conventional gentamicin protection assay followed by CFU enumeration on agar plates. One-way ANOVA with Tukey’s posttest; ns=not significant;*=p≤0.05; **= p≤0.01; ***=p≤0.001; ****=p≤0.0001

### *Y. pestis* inhibits the inflammatory capacity of hPMN-derived EVs

As EVs can also bolster cellular communication and immune cell activation, we next assessed the response of macrophages to hPMN-derived EVs. Following EV treatment, we observed significant polarization of hMDMs treated with EVs from T3E-infected cells towards an M1- phenotype, which was similar to that of cells treated with IFNγ (Figure 6b). However, hMDMs treated with EVs from Yp-infected cells did not display significant expression of M1 markers and appeared more similar to hMDMs treated with EVs from UI hPMNs. Moreover, hMDMs treated with EVs from T3E-infected hPMNs were better able to kill *Y. pestis* than cells treated with EVs from Yp-infected cells or left untreated (Figure 6c, p≤0.001). Together, these data suggest that *Y. pestis* actively inhibits the packaging of factors into EVs required to potentiate their immune stimulatory properties.

### Yop effectors act cooperatively to suppress EV protein packaging

Based on our proteomic analysis, manipulation of protein packaging into hPMN-derived EVs is dependent on the *Y. pestis* secretion of the Yop effectors. To determine the contribution of individual Yop effectors on EV protein packaging, we employed a library of *Y. pes*tis mutants that express only one Yop effector at a time, allowing us to investigate the role of each effector independent of potential functional redundancy [29, 50]. hPMNs were infected with *Y. pestis* strains from this library and changes in protein concentration were measured as an indicator of changes in EV biogenesis. Compared to EVs isolated from T3E-infected hPMNs, the protein concentrations from EVs isolated from hPMNs infected with *Y. pestis* strains expressing YopE, YopH, or YopK were significantly lower (Figure 7a; p<0.05), while no significant changes in protein concentrations were observed in EVs isolated from cells infected with *Y. pestis* expressing only YpkA, YopJ, YopM, or YopT (Figure S2). However, protein concentrations from YopE, YopH, or YopK samples remained significantly higher than cells treated with EVs from Yp-infected hPMNs (Figure 7a), indicating that single Yop effectors are not sufficient to completely inhibit EV biogenesis by hPMNs. Furthermore, co-infections with two strains expressing YopE, YopH, or YopK also failed to recapitulate the Yp phenotype, suggesting that that the cooperative functions of all three Yop effectors are required to suppress EV protein packaging (Figure 7a). Proteomic analysis of the EVs isolated from the infections with YopE, YopH, and YopK further indicates that each Yop independently influences the packaging of different proteins that are packaged with the EVs (Figure 7b).

**Figure 7:**
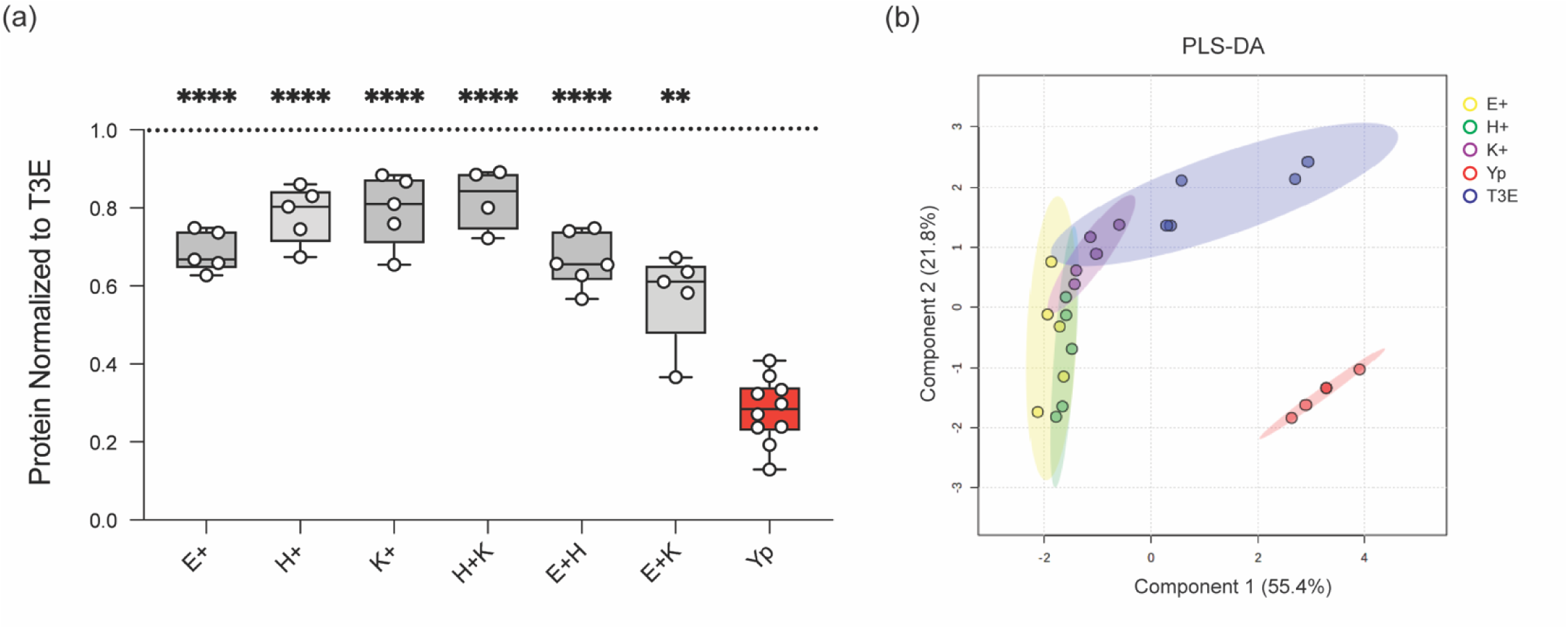
Yop effector proteins work cooperatively to manipulate EV protein packaging. hPMNs were infected with WT *Y. pestis* (Yp) or with a *Y. pestis* mutant expressing only YopE (E+), YopH (H+), or YopK (K+), or mixed at a 1:1 ratio for YopE and YopH (E+H), YopH and YopK (H+K), YopE and YopK (E+K). (a) Total protein in each EV sample was quantified and normalized to the T3E control to account for donor variability. One-way ANOVA with Dunnett’s multiple comparisons test to Yp with Geisser-Greenhouse correction; **= p≤0.01; ****=p≤0.0001 (n=5-10). (b) PLS-DA plot depicting discriminant analysis of EVs from one-dimensional reverse phase liquid chromatography tandem mass spectrometry (n=5).

**Table 1:** Bacterial Strains.

| Table 1: Bacterial Strains |  |  |
| --- | --- | --- |
| Strain | Genotype | References |
| *Y. pestis* | KIM1001, pgm<sup>-</sup>, pMT1<sup>+</sup>, pPCP1<sup>+</sup>, pCD1<sup>+</sup>, PML001<sup>+</sup> | [50] |
| *Y. pestis* T3E | KIM1001 pgm<sup>-</sup>, pMT1<sup>+</sup>, pPCP1<sup>+</sup>, pCD1<sup>+</sup> (yopH<sup>Δ3-467</sup> yopE<sup>Δ40-197</sup> yopK<sup>Δ4-181</sup> yopM<sup>Δ3-408</sup> ypkA<sup>Δ3-731</sup> yopJ<sup>Δ4-288</sup> yopT<sup>Δ3-320</sup>), pML001<sup>+</sup> | [50] |
| *Y. pestis* T3E +ypkA | KIM1001 pgm<sup>-</sup>, pMT1<sup>+</sup>, pPCP1<sup>+</sup>, pCD1<sup>+</sup> (yopH<sup>Δ3-467</sup> yopE<sup>Δ40-197</sup> yopK<sup>Δ4-181</sup> yopM<sup>Δ3-408</sup> yopJ<sup>Δ4-288</sup> yopT<sup>Δ3-320</sup>), pML001<sup>+</sup> | [50] |
| *Y. pestis* T3E +yopE | KIM1001 pgm<sup>-</sup>, pMT1<sup>+</sup>, pPCP1<sup>+</sup>, pCD1<sup>+</sup> (yopH<sup>Δ3-467</sup> yopK<sup>Δ4-181</sup> yopM<sup>Δ3-408</sup> ypkA<sup>Δ3-731</sup> yopJ<sup>Δ4-288</sup> yopT<sup>Δ3-320</sup>), pML001<sup>+</sup> | [50] |
| *Y. pestis* T3E +yopH | KIM1001 pgm<sup>-</sup>, pMT1<sup>+</sup>, pPCP1<sup>+</sup>, pCD1<sup>+</sup> (yopE<sup>Δ40-197</sup> yopK<sup>Δ4-181</sup> yopM<sup>Δ3-408</sup> ypkA<sup>Δ3-731</sup> yopJ<sup>Δ4-288</sup> yopT<sup>Δ3-320</sup>), pML001<sup>+</sup> | [50] |
| *Y. pestis* T3E +yopJ | KIM1001 pgm<sup>-</sup>, pMT1<sup>+</sup>, pPCP1<sup>+</sup>, pCD1<sup>+</sup> (yopH<sup>Δ3-467</sup> yopE<sup>Δ40-197</sup> yopK<sup>Δ4-181</sup> yopM<sup>Δ3-408</sup> ypkA<sup>Δ3-731</sup> yopT<sup>Δ3-320</sup>), pML001<sup>+</sup> | [10, 50] |
| *Y. pestis* T3E +yopK | KIM1001 pgm<sup>-</sup>, pMT1<sup>+</sup>, pPCP1<sup>+</sup>, pCD1<sup>+</sup> (yopH<sup>Δ3-467</sup> yopE<sup>Δ40-197</sup> yopM<sup>Δ3-408</sup> ypkA<sup>Δ3-731</sup> yopJ<sup>Δ4-288</sup> yopT<sup>Δ3-320</sup>), pML001<sup>+</sup> | [50] |
| *Y. pestis* T3E +yopM | KIM1001 pgm<sup>-</sup>, pMT1<sup>+</sup>, pPCP1<sup>+</sup>, pCD1<sup>+</sup> (yopH<sup>Δ3-467</sup> yopE<sup>Δ40-197</sup> yopK<sup>Δ4-181</sup> ypkA<sup>Δ3-731</sup> yopJ<sup>Δ4-288</sup> yopT<sup>Δ3-320</sup>), pML001<sup>+</sup> | [50] |
| *Y. pestis* T3E +yopT | KIM1001 pgm<sup>-</sup>, pMT1<sup>+</sup>, pPCP1<sup>+</sup>, pCD1<sup>+</sup> (yopH<sup>Δ3-467</sup> yopE<sup>Δ40-197</sup> yopK<sup>Δ4-181</sup> yopM<sup>Δ3-408</sup> ypkA<sup>Δ3-731</sup> yopJ<sup>Δ4-288</sup>), pML001<sup>+</sup> | [50] |

## Discussion

*Y. pestis* actively manipulates how PMNs respond during plague [19]. Through the action of the Yop effector proteins, *Y. pestis* inhibits phagocytosis, ROS production, degranulation, and the release of inflammatory lipids and cytokines [10, 15, 29, 50]. Together, these actions significantly impair the antimicrobial activity of these phagocytes and delay inflammation necessary to control the infection. Here we demonstrate that *Y. pestis* also alters EV biogenesis by hPMNs. Considering that EVs are key mediators of intracellular communication, these changes in EV biogenesis by Yop intoxication represent another mechanism by which the bacteria evade the immune system and delay inflammation to establish a non-inflammatory environment during infection. Defining how *Y. pestis* disrupts EV assembly will further help the field understand the pathogenesis of this bacterium and provide a new tool to investigate the molecular mechanisms that govern EV biogenesis in PMNs during bacterial infection.

In the current study, we defined the EVs produced in response to *Y. pestis* and to a mutant lacking the genes encoding the Yop effector proteins, allowing us to identify *Y. pestis*-specific hPMN responses and the impact of the Yop effectors on EV biogenesis. Several studies with other bacteria have demonstrated that EV production by PMNs increases in response to infection [59–61], but NTA analysis showed that EV production by hPMNs did not increase in response to *Y. pestis* (Figure 1a). However, we observed significant increases in EV numbers during T3E infection, indicating that *Y. pestis* actively inhibits EV biogenesis or release via the action of the Yop effectors. To our knowledge, this is the first example of a bacterial pathogen actively inhibiting EV release by host cells. While we have identified the Yop effectors responsible for this phenomenon, we have yet to identify the specific molecular mechanisms responsible for this inhibition. Moreover, because we have not differentiated small and large EVs in our samples, it remains to be determined if this is a global inhibition of all EVs, or perhaps inhibition of a specific pathway (i.e., plasma membrane budding of large EVs vs. multivesicular body production of small EVs). While clear differentiation of these populations is difficult, the Yop effector mutants provide us with tools to better understand the mechanisms of EV biogenesis during infection.

Directly correlating with the differences in the number of EVs released, we also observed significantly more protein in EVs isolated from hPMNs infected with the T3E mutant compared to our other samples (Figure 2a). More interesting is the clear differences in the proteins that were packaged into EVs during Yp and T3E infections. In general, these proteins could be grouped into three categories: those overrepresented in the UI samples, those enriched in both infections, and those overrepresented in T3E infected hPMNs. These differences indicate that in addition to blocking EV release, *Y. pestis* also actively alters the host proteins selectively packaged into the EVs.

One group of proteins significantly enriched in the UI samples compared to the samples from bacterial infections were histone proteins. While histones are often considered common contaminants of EV isolation, the significant differences in enrichment we observed between our three conditions, especially the relative absence in the T3E samples, suggest a more nuanced response and perhaps intentional EV packaging. Specific EV packaging of histones in cells undergoing cellular stress has been suggested by previous studies showing dramatic changes in histone colocalization within the multivesicular body, where small EVs are produced [62].

While the implications of differential histone packaging in EVs during infection has yet to be defined, these data suggest active changes in nucleosome assembly and histone localization during infection, which is altered by Yop injection. Such changes could have implications on neutrophil gene expression or induction of neutrophil extracellular traps, both which merit further consideration in the context of plague.

In contrast to histones, we observed several classes of proteins that were overrepresented in EVs from infected hPMNs, regardless of the *Y. pestis* strain. Among these were proteins associated with the extracellular matrix (ECM) and nutritional immunity (Figures 4c and 4d). The packaging of ECM related-proteins within EVs is not uncommon, with Halawani et al. suggesting that ECM-associate proteins comprise ∼12% of the EV proteome on average [63]. In our study, the ECM-related proteins that were enriched included key proteases that can directly degrade ECM and promote inflammation, including matrix metalloproteinases MMP8, MMP9, and neutrophil elastase (ELANE). Recently, Nudelman et al. reported a similar enrichment for MMP9 in EVs produced by macrophages in response to *Salmonella enterica* Typhimurium [64]. These EVs, which they referred to as proteolytic EVs, appeared to be released in response to bacteria and enhanced macrophage migration through ECM, suggesting that the presence in these proteases in EVs released by infected PMNs may also contribute to migration or invasion into infected tissues.

Proteins associated with nutritional immunity were among the most abundant peptides identified in EVs from infected cells. Nutritional immunity is an arm of the innate immune system that restricts access to essential metals (e.g., iron, zinc, and manganese) from invading pathogens by the release of host proteins that scavenge these metals. Among these, Lactoferrin (LTF), Lipocalin (LCN2), and Haptoglobin (HP) restrict microbial access to iron by direct sequestration of iron, bacterial produced siderophores, or heme by binding to hemoglobin, respectively [65]. Like the ECM-related proteins mentioned above, these nutritional immunity proteins are stored within the specific granules of PMNs and typically released in response to infection during degranulation. While it is possible that EVs may acquire these proteins post-biogenesis if they are released after degranulation, Yp effectively inhibits degranulation by PMNs [28, 29], suggesting that crosstalk between the EV biogenesis pathway and PMN granules is more likely contributing to the packaging of granule proteins prior to release. Unlike the majority of the nutritional immunity proteins that were associated with EVs isolated from infected hPMNs, calprotectin (S100A8/S100A9) is not stored within granules but is found within the cytosol of the cell [66], indicating additional mechanisms of EV protein packaging in PMNs. Bode *et al*. suggested that cell stimulation can promote association of calprotectin with Annexin 6, resulting in increased localization at the plasma membrane [67]. Such active colocalization at the membrane could increase the incorporation of calprotectin into large EVs via plasma membrane budding. Cytosolic localization of calprotectin within PMNs has also raised questions regarding how this important metal sequestration protein is rapidly released during infection, as it would not occur via conventional degranulation. Other studies have suggested release during NETosis [68] or via inflammasome-mediated gasdermin pore formation [69], but our data suggest that calprotectin may also be released from the cell via EVs in response to infection. Additional studies are required to assess the overall contribution of EVs to the total calprotectin release by neutrophils and localized metal sequestration.

The final trend we observed was the enrichment of specific proteins in EVs following T3E infection that were absent or significantly lower in EVs from Yp-infected cells. These represent proteins that appear to be specifically excluded during EV biogenesis by the activity of the Yop effector proteins. Among this group were numerous Annexins, which were present in EVs from UI and T3E-elicited EVs but significantly underrepresented in EVs from Yp-infected hPMNs (Figure 5d). Annexins have well-established roles in vesicular trafficking, especially membrane trafficking, and in EV biogenesis [70–72]. They have been linked to MVB fusion to the plasma membrane, membrane budding, cargo sorting, and the timing of EV release by cells [70–72]. Annexin-mediated trafficking is responsive to changes in calcium signaling within the cell, with calcium binding leading to trafficking to and interactions with phospholipids within membranes [72]. A key effect of YopH on the PMNs is the inhibition of calcium flux [18], which would indirectly inhibit Annexin activation and trafficking, and likely contributing to the impact of YopH on altering EV biogenesis.

In addition to the Annexins, the other category of proteins significantly enriched in the T3E- elicited EVs were several antimicrobial proteins. Similar to the nutritional immunity and ECM-associated proteins, these proteins are found within the granules but associated with the azurophilic granules at a much higher frequency than the former groups of proteins. Exclusion of these antimicrobial proteins suggest that *Y. pestis* may specifically alter interactions between EVs with the azurophilic granules during EV biogenesis, effectively limiting incorporation of cargo from these compartments into EVs [57]. The enrichment of these proteins, especially myeloperoxidase (MPO), which was the second most abundant protein recovered from T3E- elicited EVs, are likely responsible for the potent bactericidal activity of these EVs (Figure 6a). These data support previous reports that PMNs have a unique ability to generate antimicrobial EVs as part of immune response repertoire [32, 35, 58], but is blocked by *Y. pestis*.

Intercellular communication is a paramount function of EVs. We showed that EVs isolated from T3E-infected, but not Yp-infected cells, effectively influenced the activation or polarization of macrophages (Figure 6b), similar to findings of Duarte *et al*. reporting diminished hMDM activation after treatment with PMN-derived EVs that were produced during infection with *Mycobacterium tuberculosis* [73]. As PMNs represent the primary population of immune cells that interact with *Y. pestis* during colonization [7, 9], limiting the inflammatory potential of the EVs released by these cells likely contributes to the delayed inflammatory response observed during plague. While we have demonstrated dramatic changes in the proteins packaged within Yp-elicited EVs, the lipids and small RNAs that are packaged within EVs also significantly contribute to intercellular communication. For example, leukotriene B4, a potent chemoattractant and activator of immune cells, is typically rapidly produced and packaged in EVs in response to infection [43] but its synthesis is actively inhibited by *Y. pestis* [10, 29], and thus absent during EV biogenesis. We anticipate that Yp also disrupts the packaging of other lipids and small RNAs that are typically packaged into PMN-derived EVs in response to infection. Studies are ongoing to define the changes of these components during Yp and T3E infection within PMNs and the potential role of these molecules in influencing intercellular communication.

Inhibition of EV biogenesis was dependent on secretion of the Yop effectors, specifically the cooperative action of YopE, YopH, and YopK (Figure 7). While YopH is a tyrosine phosphatase that specifically targets the SKAP2/SLP-76/PRAM-1 signaling hub to inhibit PLCγ2-mediated calcium signaling [14, 74], YopE is a Rho GTPase-activating protein (GAP) that inactivates RhoA and Rac1 in neutrophils to disrupt the actin cytoskeleton [17]. Both calcium signaling and RhoA/Rac1 activity have been previously implicated in EV trafficking and uptake by recipient cells [75–78]. Moreover, we have previously shown that inhibition of degranulation also required the cooperative activities of YopE and YopH, suggesting that YopE and YopH inhibit the crosstalk between EVs and the azurophilic granules needed for the packaging of the antimicrobial proteins into the EVs. Indeed, MS analysis of EVs from hPMNs infected with YopH or YopE only strains showed reduced enrichment of MPO and other proteins selectively enriched in T3E-elicited EVs (Figure S3). Unlike YopE and YopH, YopK has not been shown to directly impact trafficking pathways in neutrophils. Instead, YopK is thought to regulate the translocation of other Yops into the cell, including the YopB and YopD translocase [79], to inhibit inflammasome activation in macrophages [80]. Inflammasome activation has been directly linked to EV production [81, 82], suggesting that limiting activation of this important response during infection also contributes to changes in the proteins that are packaged into the EVs and their downstream signaling compacity. Interestingly, this is not the first example of the cooperation between these three Yop effectors. A similar requirement of YopE, YopH, and YopK was reported for complete inhibition of caspase-4 inflammasome activation in human macrophages [83].

In conclusion, we have discovered a potential role for EVs in maintaining the non-inflammatory environment essential for plague pathogenesis. The *Y. pestis* T3SS effectors directly manipulate EV packaging, effectively blunting immune cell communication and dampening immune recognition of infection. Further, we have shown that manipulation of EV biogenesis by the Yop effector proteins limits the antimicrobial and immune modulatory capacity of neutrophil derived EVs, limiting the potential of these EVs to thwart off pathogens and relay immunologic signals. While the specific molecular mechanisms by which *Y. pestis* manipulates EV cargo require further investigation, these studies have demonstrated that *Y. pestis* can be used as a tool to further investigate the nuances of EV biogenesis and their role in responses to bacterial infection.

## Acknowledgements

This work was supported in part by funds from the NIH NIAID grants F31AI178999 (KRS), R21AI169423 (MBL, SMU), R01AI178106 (MBL), R01AI148241 (MBL), the UofL Hepatobiology and Toxicology COBRE NIGMS grant P20GM113226 (JH), the University of Louisville Kidney Disease Program and the Proteomics Technology Center (TDC, MLM) the NIGMS-funded KY INBRE P20GM103436 (KRS), and the Jewish Heritage for Excellence Fund (MBL).

The authors would like to acknowledge Dr. Julia Aebersold with the Center for Micro/Nano Technology at the University of Louisville for assistance with TEM. We would also like to acknowledge Dr. Kevin Sokoloski at the University of Louisville for helping us with biological processes pathway analysis figure generation.

## Declaration of Interest Statement

All authors report no conflicts of interest.

## Data availability statement

The acquired proteomic data files will be deposited in MassIVE (http://massive.ucsd.edu/) data repository and shared with ProteomeXchange (https://www.proteomexchange.org/) using the Project title “*Yersinia pestis* actively inhibits the production of extracellular vesicles by human neutrophils”. These data include (A) the primary data files (.RAW) for the fractionated human neutrophil proteome (B) an excel file with assembled Peaks X search results, (C) the sample key, and (D) the human reviewed canonical FASTA sequence database. These files will be released from embargo upon manuscript acceptance.

---

**Supplemental Figure S1:**
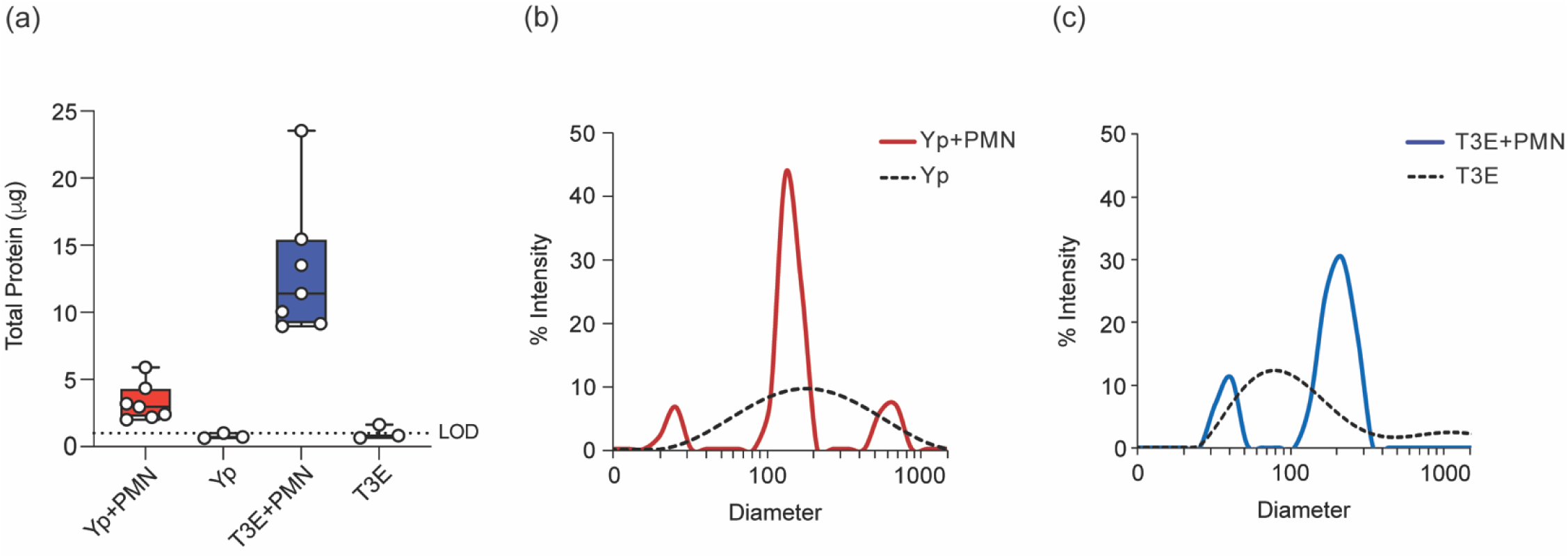
**Negligible OMV contamination within EV prep.** WT *Y. pestis* (red) or T3E *Y. pestis* (blue) was incubated with or without hPMNs for 1 h prior to EV isolation. (a) Protein quantification was performed using Protelite Fluorometric Protein Quantification Kit optimized for the Qubit fluorometer. Each point represents an independent biological replicate. (b-c) DLS analysis depicting EV profiles comparatively. Representative results of 3 independent experiments.

**Supplemental Figure S2:**
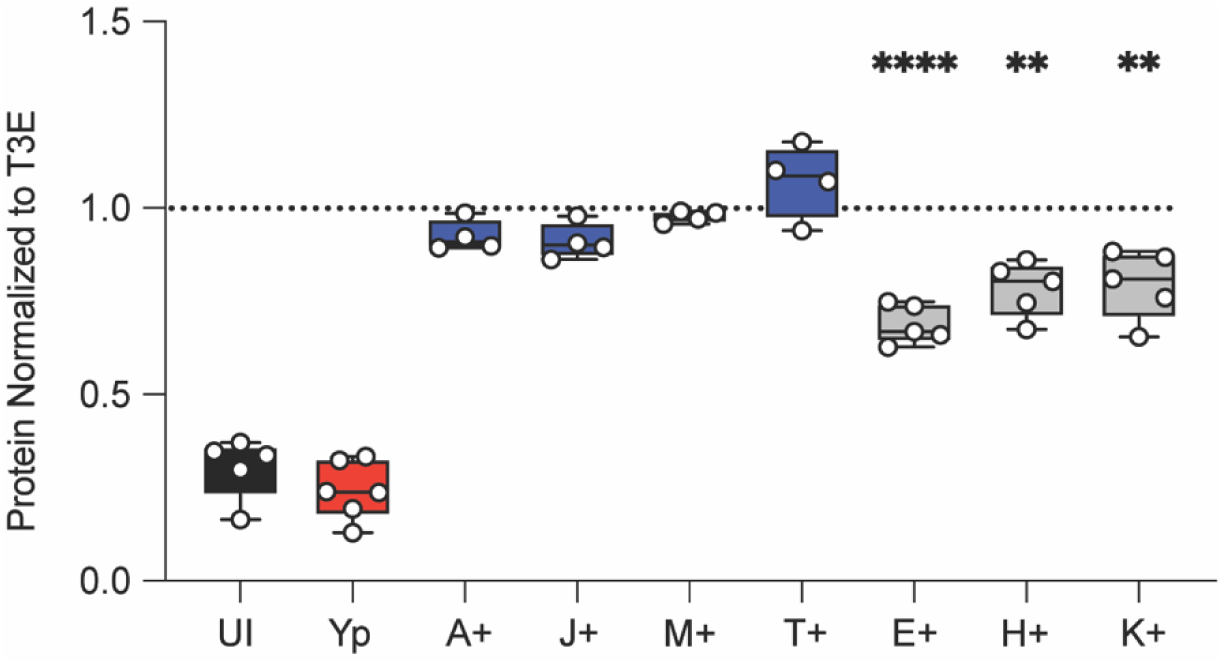
**Effect of individual Yop effectors on hPMN EVs.** EVs were isolated from uninfected hPMNs (UI) or hPMNs that were infected with WT *Y. pestis* (Yp) or with a *Y. pestis* mutant expressing only YopA (A+), YopJ (J+), YopM (M+), YopT (T+), YopE (E+), YopH (H+), or YopK (K+). Total protein in each EV sample was quantified and normalized to the T3E control to account for donor variability. Recorded statistical significance is relative to T3E. One-way ANOVA with Dunnett’s multiple comparisons test to T3E with Geisser-Greenhouse correction; **= p≤0.005; ****=p≤0.0001 (n=4-6).

**Supplemental Figure S3:**
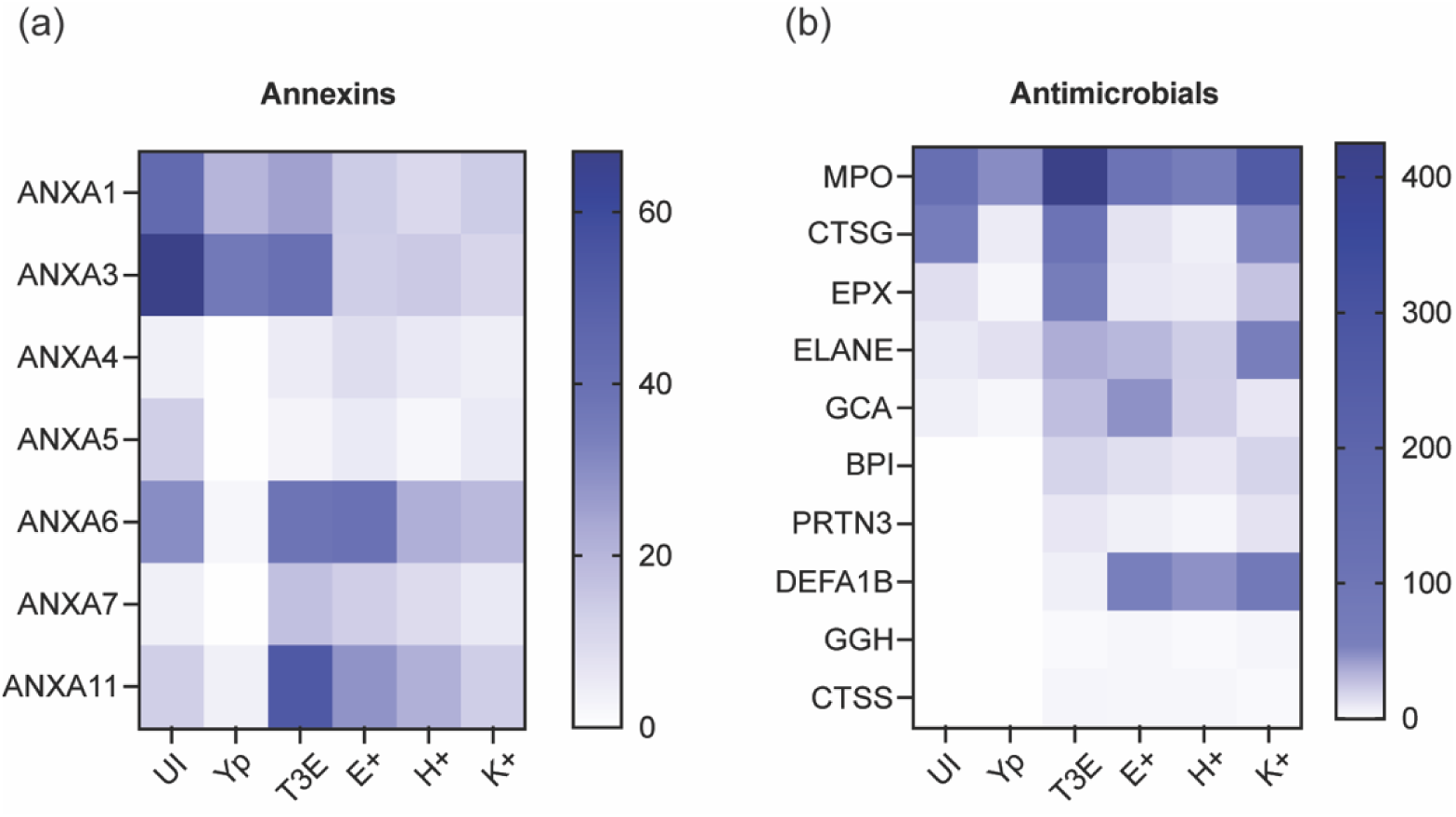
**Impact of individual Yop effectors on EV protein packaging.** EVs elicited from hPMNs infected with *Y. pestis* mutants expressing a single Yop effector were analyzed via mass spectrometry. Comparative enrichment of annexin (a) and antimicrobial proteins (b).

## Notes

http://massive.ucsd.edu/

https://www.proteomexchange.org/

### Competing Interest Statement

The authors have declared no competing interest.

